# Effects of Gallic Acid on Antioxidant Defense System in Mice with Benzene-Induced Myelotoxicity

**DOI:** 10.1101/2024.06.12.598751

**Authors:** Toba Isaac Olatoye, J.O. Adebayo

## Abstract

Benzene is known to cause myelotoxicity which impacts secondary complications mediated by oxidative stress on the heart and erythrocyte. Gallic acid is an antioxidant which has not been evaluated for protective effect against cardiovascular complications of benzene-induced myelotoxicity. Therefore, this study was carried out to evaluate the effects of gallic acid on the concentrations of selected oxidative stress biomarkers and activities of antioxidant enzymes in the erythrocyte and heart of mice with benzene-induced myelotoxicity. Thirty-six male mice were randomized into six groups of six mice each. Group A served as the normal control receiving distilled water while the remaining groups were orally administered 150 mg/kg body weight of benzene for fourteen days. Distilled water, 50 mg/kg body weight of ascorbic acid, 25, 50, and 100 mg/kg body weight of gallic acid were simultaneously administered to mice of groups B (negative control), C, D, E, and F respectively for fourteen days. Concentrations of oxidative stress biomarkers and activities of antioxidant enzymes in target tissues were then determined. Results revealed significant elevation (p<0.05) in the concentrations of malondialdehyde, nitric oxide and protein carbonyl, as well as significant decrease (p<0.05) in the concentrations of reduced glutathione, protein, and activities of catalase, glutathione peroxidase, glutathione-S-transferase and superoxide dismutase in the heart and erythrocyte of negative control compared to normal control. However, treatment with gallic acid at various doses significantly reverted (p<0.05) the observed alterations in these parameters, comparing favourably with ascorbic acid (reference drug). These results suggest that gallic acid protected the heart and erythrocyte against lipid and protein oxidation, and enhanced antioxidant defense system of mice with benzene-induced myelotoxicity.

## Introduction

Myelotoxicity, also known as myelosuppression or bone marrow suppression, is the decrease in production of cells responsible for providing immunity (leukocytes), carrying oxygen (erythrocytes), and those responsible for normal blood clotting (thrombocytes) (Parent-Massin *et al*., 2010). Myelotoxicity is a serious side effect of chemotherapy and certain drugs affecting the immune system such as azathioprine (Robert *et al*., 2000). Clinical signs of bone marrow suppression include bleeding, caused by a reduction in platelet counts; anemia, which leads to fatigue and altered cardiovascular or respiratory parameters; and a heightened susceptibility to various infections due to leukocytes repression (Williams *et al*., 1997).

A variety of pharmaceuticals and industrial chemicals can cause partial or complete bone marrow suppression, which may be reversible or permanent depending on the chemical agent and extent of exposure (Hancock *et al*., 1984). Benzene is the best-known bone marrow suppressant among industrial chemicals. It was used decades ago to inhibit the uncontrollable production of leukemia cells (Kippen *et al*., 1988). Today, cancer chemotherapeutics especially the alkylating agents are the most frequently encountered causes of myelotoxicity (Robert *et al*., 2000).

In humans, benzene-induced myelotoxicity usually occurs as a result of occupational exposure to benzene (Cooper and Snyder, 1988; Yin *et al*., 1987). Several mechanisms have been proposed to explain benzene-induced myelotoxicity but, despite extensive research, the mechanism by which benzene induces myelotoxicity is still not clear. However, studies have shown that benzene metabolites are the primary causes of oxidative stress experienced during myelotoxicity (Rickert *et al*., 1979; Irons *et al*., 1982; Powley *et al*., 2000). Benzene is metabolized by cytochrome P450 2E1 (CYP2E1), one of the many isozymes of hepatic cytochrome P450 mixed function oxidases (Irons *et al*., 1982; Lovern *et al*., 1997; Powley *et al*., 2000). Benzene oxide, the first intermediate in CYP 2E1-mediated metabolism, is converted into a number of metabolites including phenol, hydroquinone, benzoquinone, catechol, and muconic acid/muconaldehyde. In the bone marrow, myeloperoxidase further oxidizes these phenolic metabolites of benzene to form free radicals capable of damaging the bone marrow (Sawahata *et al*., 1985; Kipen *et al*., 1988). The damage inflicted in bone marrow by these toxic radicals sets the stage for numerous secondary complications that frequently emerge. Oxidative stress has been implicated as either the primary cause of pathologies such as atherosclerosis, paraquat poisoning, radiation-induced pneumonitis and fibrosis, or as secondary contributor to disease progression including chronic obstructive pulmonary disease (COPD), cancers, type II diabetes, hypertension, cardiovascular and Alzheimer’s disease (Henry and Zhang, 2021).

However, since the role of oxidative stress in many diseases remains incompletely understood, and given that oxidative stress is primarily responsible for the pathology of bone marrow suppression and its consequent cardiovascular complications, in the presence of an impaired antioxidant defense system, therefore it is crucial to investigate the role of gallic acid, an exogenous antioxidant derived from the diet, which has not been previously studied in this context, in assisting the cell’s first line of defense against secondary complications mediated by oxidative stress in heart and erythrocyte of mice exposed to benzene.

## Materials and Methods

### Chemicals and reagents

Benzene, gallic acid, ascorbic acid, and sesame oil were purchased from Tokyo Chemical Industry Co. Ltd, Tokyo, Japan. Assay kits used for antioxidant enzymes and oxidative stress biomarkers assay were products of Randox Laboratory Ltd, Ardmore,Co., Antrim, UK. Other chemicals used were obtained commercially and of analytical grade.

### Experimental animals

Thirty-six (36) male mice, about 4 months old and average weight 27 ± 2 g were used for the experiment. The mice were obtained from the Animal Holding Unit of the Department of Biochemistry, University of Ilorin, Ilorin, Nigeria, and were acclimatized for 14 days under standard housing conditions (temperature, 25 ± 2 □C; humidity, 50-70%; 12 hrs. light/dark cycle). They were allowed free access to clean water and standard diet *ad libitum*.

### Ethical clearance

Ethical clearance on the use of laboratory animals was issued by Ethics Committee of University of Ilorin, and the researcher adhered strictly to the Principles of Laboratory Animal Care (NIH Publication, No. 85-23).

### Experimental study design

The mice were randomized into six groups of 6 animals each. The grouping was done as follows:

Group A (Normal Control): 0.2 ml/20 g distilled water + 0.2 ml/20 g sesame oil.

Group B (Negative Control): 150 mg/kg body weight benzene + 0.2 ml/20 g distilled water.

Group C: 150 mg/kg body weight benzene + 50 mg/kg body weight ascorbic acid (reference drug).

Group D: 150 mg/kg body weight benzene + 25 mg/kg body weight gallic acid.

Group E: 150 mg/kg body weight benzene + 50 mg/kg body weight gallic acid.

Group F: 150 mg/kg body weight benzene + 100 mg/kg body weight gallic acid.

Doses of benzene, gallic acid, and ascorbic acid were sourced from literature (You and Park 2010; Inoue and Hirabayashi 2010;Wei *et al*., 2017).

### Induction of myelotoxicity

Myelotoxicity was induced in mice via oral administration of benzene after it was diluted in sesame oil, in a single dose of 150 mg/kg body weight, once daily, 6 days/week, for 14 days. Ascorbic acid and gallic acid were also administered the same way simultaneously with benzene throughout the duration.

### Blood sample collection

At the end of 14 days (24 hrs. after the last treatment), the animals were anaesthetized with diethyl ether and venous blood was collected into EDTA bottles.

### Blood sample preparation

The blood was centrifuged (4000 rpm for 10 min, 4 °C) and plasma was separated. After removing the buffy coat, red blood cells (RBCs) were washed in 4 ml of phosphate buffer (100 mM, pH 7.4) twice. Following the second washing, supernatants were removed and packed RBCs were obtained. All samples were stored at -80 °C until needed for further analyses.

### Red blood cell lysis

10 ml of 10x RBC lysis buffer per 1 ml of blood was added and incubation was done for 10-15 min at room temperature. Turbidity was observed until it disappeared. Centrifugation of the lysate at 3500 rpm for 5 mins was done, the supernatant was pipetted out and frozen at -80 ^o^C until needed for various biochemical analyses of this study.

### Preparation of tissue homogenate

The mice were quickly dissected and their hearts excised, cleansed of blood stains with cotton wool, weighed, and homogenized separately in ice-cold 0.25 M sucrose solution (1:5, w/v). The homogenates were centrifuged at 3500 rpm for 5 min to obtain the supernatants which were pipetted out and frozen at -80 ^o^C until needed for further analyses.

## Biochemical analysis

### Determination of oxidative stress markers

#### Nitric oxide (NO)

Nitric oxide concentration of heart homogenates and erythrocyte lysates was estimated using Griess reaction (Guevara *et al*., 1998).

#### Principle

The principle of this assay is based on the enzymatic conversion of nitrate to nitrite by nitrate reductase. Nitric oxide level was expressed as mg/dl protein.

#### Procedure

Sodium nitroprusside (20 µl) was added to 50 µl of the sample and incubated at 37^°^ C for 30 min. Then, 100 µl of Griess reagent was added to the mixture. The absorbance reading was taken at 550 nm.

#### Malondialdehyde (MDA)

MDA of heart and erythrocyte, a marker of lipid peroxidation was determined using the method of Hunter (1963) as modified by Gutteridge and Wilkins (1982). The MDA level was expressed as µmol/ml protein.

#### Principle

MDA, a product of lipid peroxidation, when heated with 2-thiobarbituric acid (TBA) under acid conditions forms a pink-coloured product which has a maximum absorbance of 532 nm.

#### Procedure

The homogenates of heart and erythrocyte lysates were each supplemented with 1 g of TBA in 100 ml of 0.2% NaOH and 3 ml of glacial acetic acid, thoroughly mixed and incubated in boiling water bath for 15 minutes, after cooling, they were centrifuged. Absorbance was read at 532 nm and the results expressed as µmol/ml protein.

#### Protein Carbonyl (PCO)

As a hallmark of protein oxidation, total PCO concentration of heart and erythrocyte was determined by a spectrophotometric method described by Graziano *et al*., 2015.

#### Principle

This assay is based on 2,4-dinitrophenylhydrazine (DNPH) also known as Brady’s reagent, which is a specific probe that is able to react with PCO leading to formation of protein conjugated dintrophenylhydrazones (DNP). PCO concentration was expressed as µmol/ml/mg protein.

#### Procedure

Dinitrophenylhydrazine (DNPH) (100 µl of 10 mM) was added to 100 µl of sample in an Eppendorf tube and incubated in the dark for 1 h. Then, 100 µl of Trichloroacetic acid (TCA) was added and centrifuged at 8000 rpm for 3 min. The supernatant was discarded, and the pellet was washed with ethyl acetate and ethanol in 1:1 (500 µl) till it dissolved. Centrifugation and washing were done again and the process repeated thrice. The pellet was dissolved in 500 µl of 6 M guanidine hydrochloride and incubated for 10 min at 37 ^o^C, followed by centrifugation at 8000 rpm for 3 min. A 100 µl of the supernatant was taken and the absorbance read at 370 nm.

#### Protein concentration

Protein concentration was assayed for, using Bradford method described by Bradford, 1976 and expressed as mg/dl protein.

#### Principle

The principle of this assay is that the binding of protein molecules to Coomassie dye under acidic conditions results in a color change from brown to blue.

#### Procedure

The protein sample (30 μl) in triplicate was added to 30 μl of protein preparation buffer (0.01 M, pH 7.4), followed by the addition of 1.5 ml of Bradford reagent. Incubation was carried out at room temperature for 5 min and the absorbance taken at 595 nm.

## Determination of enzymatic antioxidant

### Superoxide dismutase (SOD)

A method originally described by Misra and Fridovich, 1972 was employed.

### Principle

The principle of this assay is based on the ability of SOD to inhibit the autoxidation of epinephrine. One SOD unit corresponds to the enzyme required to inhibit half of the oxidation of epinephrine and was expressed as U/mg protein.

### Procedure

The homogenates of heart and erythrocyte lysates (2 ml each) were supplemented with 2.5 ml of carbonate buffer (0.09 M, pH 10.2), followed by equilibration at room temperature; 0.3 ml of 0.3 nM epinephrine solution was then added to the reference and test solution, followed by mixing and reading of absorbance at 380 nm.

### Catalase (CAT)

Activities of catalase in heart and erythrocytes were determined according to the procedure of Sinha (1972).

### Principle

This principle is based on the reduction of dichromate in acetic acid to chromic acetate when heated in the presence of H_2_O_2_, with the formation of perchromic acid as an unstable intermediate. The chromic acetate so produced is measured spectrophotometrically by reading the absorbance at 380 nm within 0-90 seconds against distilled water. One unit of CAT activity is defined as the amount of enzyme required to decompose 1 mmol of H_2_O_2_ and was expressed as mmol H_2_O_2_ utilized/min/mg protein.

### Procedure

The assay mixture contained 0.5 ml of H_2_0_2_, 1.0 ml of phosphate buffer (0.01 M, pH 7.4) and 0.4 ml water. Sample (0.2 ml) was added to initiate the reaction. Dichromate/acetic acid reagent (2.0 ml) was added after 0, 30, 60, 90 seconds of incubation. To the control tube, homogenates (2 ml each) were added after the addition of the acid reagent. The tubes were then heated for 10 min and the colour developed was read at 380 nm.

### Glutathione Peroxidase (GPx)

GPx activity in each homogenate and lysate was determined in triplicate using a method described by Paglia and Valentine (1967).

### Principle

The assay is based upon the GPx mediated oxidation of GSH during reduction of cumene hydroperoxide substrate. In the presence of glutathione reductase and NADPH, the oxidized glutathione is immediately converted to the reduced form with a concomitant oxidation of NADPH. The diluting agent reduces any GSSG present in the hemolysates to GSH because the cyanide in the Drabkin reagent rapidly inactivates GSSG. A unit of GPx activity in each sample is defined as 1 nmole NADPH oxidized/min. The activity of GPx was expressed in terms of mmol of NADPH oxidized/min/mg protein.

### Procedure

Homogenate (5 ml) was diluted with 1 ml Ransel diluting agent and incubated for 5 min, followed by addition of 1 ml Drabkin reagent and mixing (total dilution of homogenate before assay, 1:82). In the actual assay, 0.1 ml of the diluted homogenate was combined with a solution containing 1 unit of the reductase and 0.1 ml of 0.2 mM NADPH. Upon addition of 40 µl 1.5 mM cumene hydroperoxide solution, the absorbance was monitored at 340 nm over a 3 min period in a spectrophotometer to determine the rate of conversion of the NADPH to NADP^+^.

### Glutathione-S-tranferase (GST)

The determination of glutathioneS-transferase activity was performed according to Habig *et al*., (1974) using 1-chloro-2,4-dinitrobenzene (CNDB) as a substrate. GST activity was expressed as mmol of CNDB formed/min/mg protein.

### Principle

GST catalyzes the conjugation of L-glutathione to CDNB. The reaction product, GS-DNB conjugate, absorbs light at 340 nm. The rate of increase in the absorption is directly proportional to the GST activity in the sample.

### Procedure

Briefly, 180 µl GST buffer (9.8 ml of phosphate buffer saline), 100 µl GSH and 100 µl CDNB were added to 20µl sample. The absorbance was read at 340 nm for 5 min every 1 min.

## Determination of non-enzymatic antioxidant

### Reduced glutathione (GSH)

The glutathione concentration of heart homogenates and erythrocyte lysates was estimated according to the method described by Sedlak and Lindsay (1968).

### Principle

The assay is based on the glutathione recycling system by 5,5-dithiobis-2-nitrobenzoic acid (DTNB) and glutathione reductase. DTNB and glutathione (GSH) react to generate 2-nitro-5-thiobenzoic acid which has yellow color. The absorbance was read at 412 nm using spectrophotometer. Total GSH content was expressed as µmole/mg of protein.

### Procedure

To 1 ml of the sample suspension (1 mg protein/ml), 1 ml of 10% Trichloroacetate (TCA) containing 1 mM EDTA was added. The protein precipitate was separated by highspeed centrifugation at 2500 rpm for 10 min. About 1 ml of supernatant was treated with 5 ml of Ellmans reagent and 3 ml of phosphate buffer (0.2 M, pH 7.4). The absorbance was read at 412 nm using spectrophotometer.

### Statistical analysis

All data were expressed as the means of six determinations ± standard error of mean (S.E.M). Statistical evaluation of data was performed using one-way analysis of variance, followed by Tukey’s test as posthoc test using Graphpad Prism (Graphpad Software Inc., California, USA) version 8.0. Significant levels were tested at p < 0.05.

## Results

### Malondialdehyde (MDA)

Benzene caused significant increase in erythrocyte and heart MDA concentration of negative control compared to normal control. However, gallic acid at all doses significantly reverted the observed increase in heart MDA concentration of negative control to the range of normal control, comparing favourably with ascorbic acid. Gallic acid only at the highest dose significantly reverted the observed increase in erythrocyte MDA concentration of negative control to the range of normal control while other doses and ascorbic acid did not cause any significant change compared to negative control (Figure 1).

**Figure 1.**
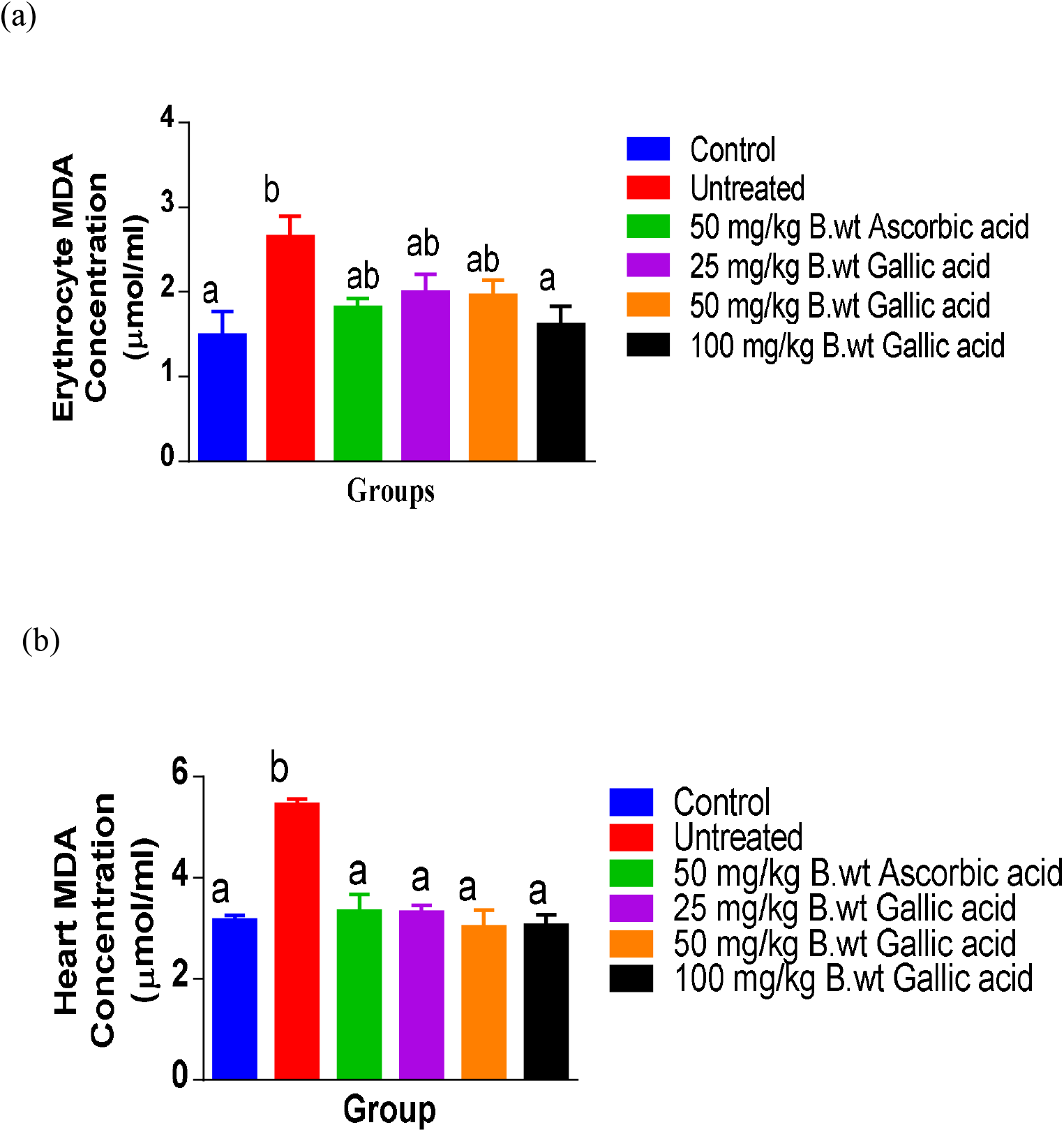
Effects of gallic acid on erythrocyte (a) and heart (b) malondialdehyde (MDA) concentration of mice with benzene-induced myelotoxicity. Values are means ± SEM of 6 replicates. Values for heart/erythrocyte with different superscripts are significantly different at p<0.05.

### Protein carbonyl (PCO)

Benzene caused significant increase in erythrocyte and heart PCO concentrations of negative controls compared to normal controls. However, gallic acid at all doses significantly reverted the observed increase in negative controls to the range of positive controls, comparing favourably well with ascorbic acid (Figure 2).

**Figure 2.**
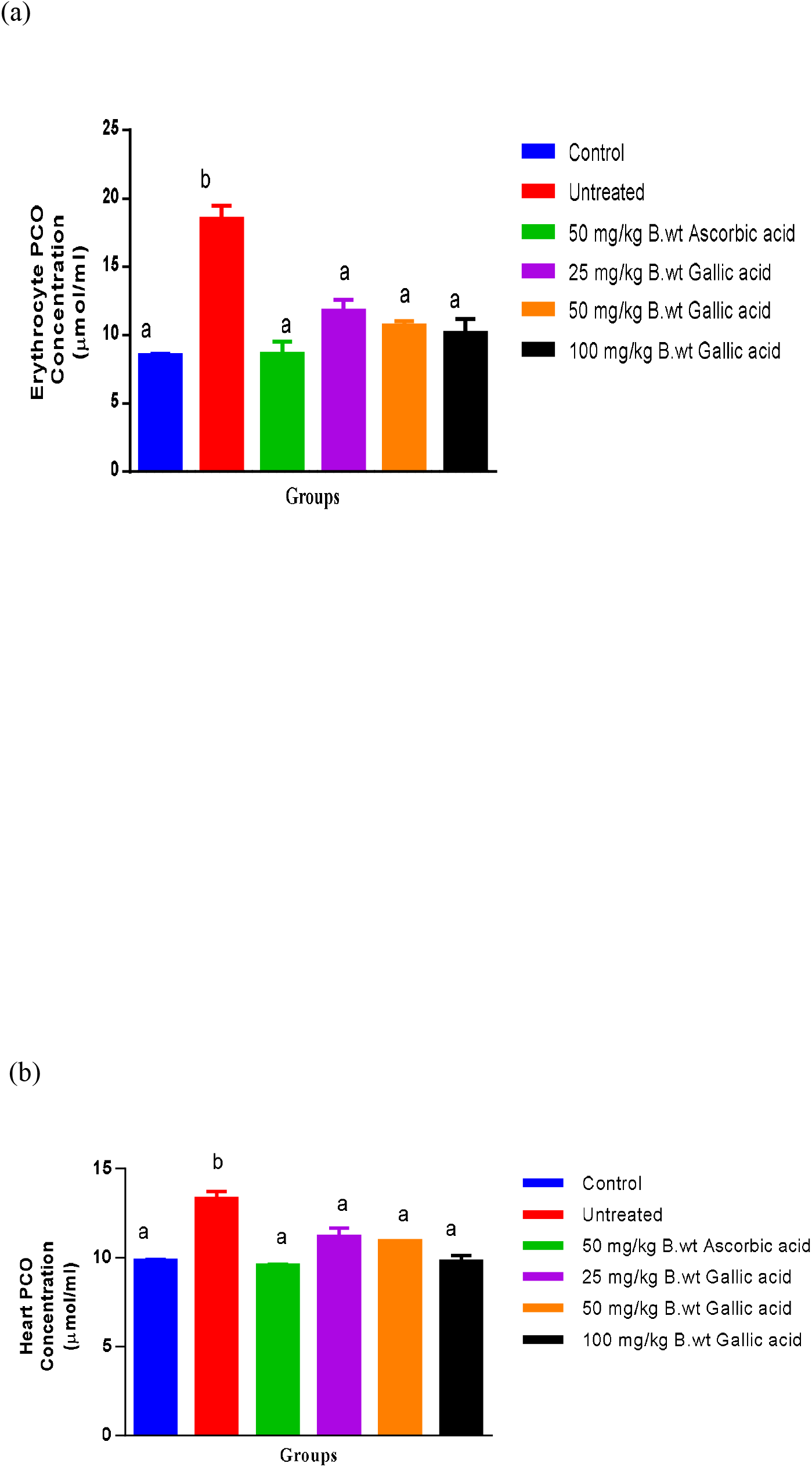
Effects of gallic acid on erythrocyte (a) and heart (b) protein carbonyl (PCO) concentration of mice with benzene-induced myelotoxicity. Values are means ± SEM of 6 replicates. Values for heart/erythrocyte with different superscripts are significantly different at p<0.05.

### Nitric oxide (NO)

Benzene caused significant increase in erythrocyte and heart NO concentrations of negative control compared to normal control. However, gallic acid at all doses significantly reverted the observed increase in erythrocyte NO concentration to the range of normal control while only doses higher than 25 mg/kg body weight significantly reverted the observed increase in heart NO concentration to the range of normal control, comparing favorably well with ascorbic acid (Figure 3).

**Fig 3.**
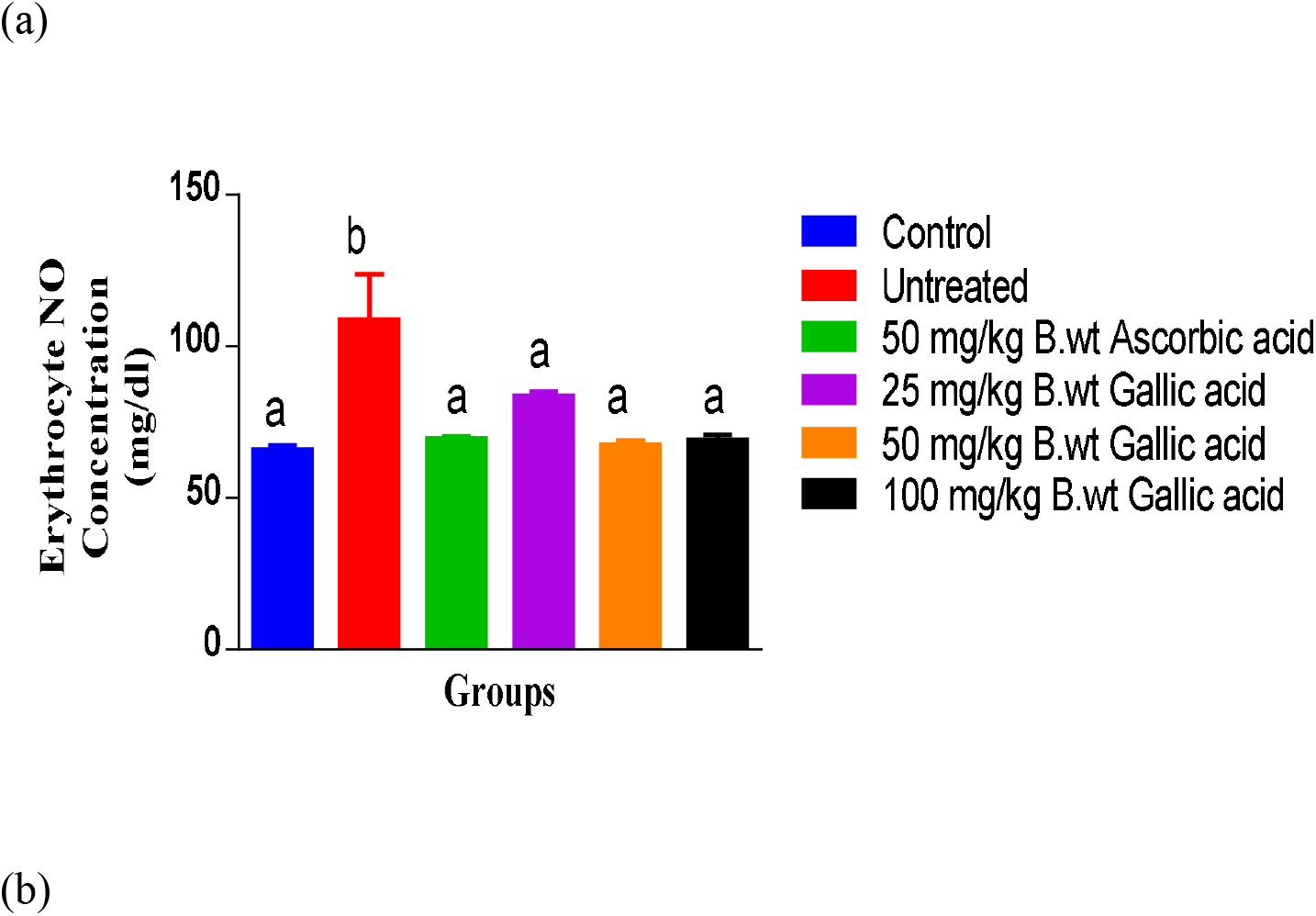
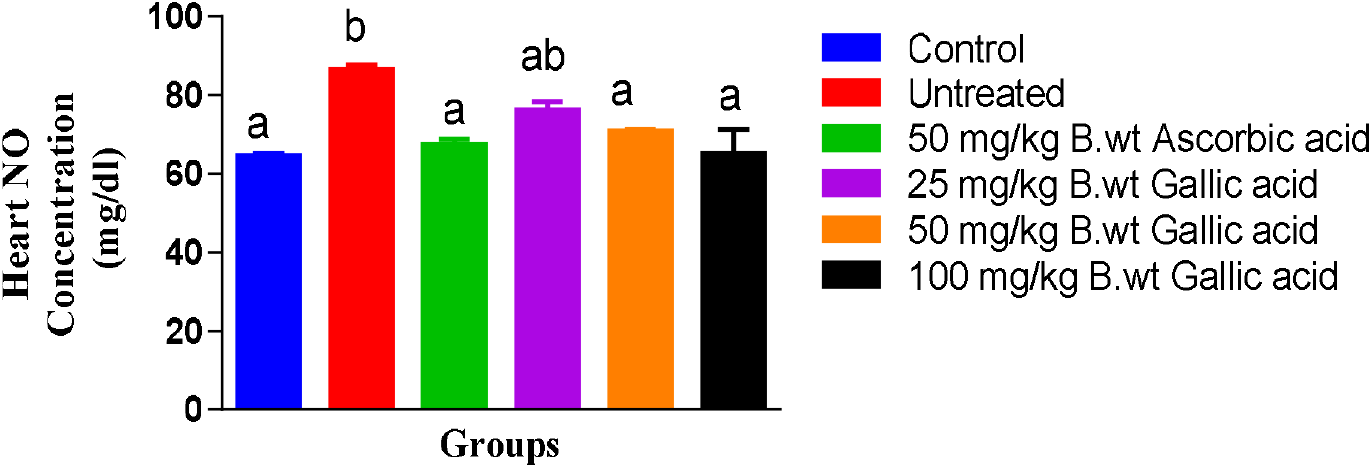
Effects of gallic acid on erythrocyte (a) and heart (b) nitric oxide (NO) concentration of mice with benzene-induced myelotoxicity. Values are means ± SEM of 6 replicates. Values for heart/erythrocyte with different superscripts are significantly different at p<0.05.

### Superoxide dismutase (SOD)

Benzene caused significant reduction in erythrocyte and heart SOD activities of negative control compared to normal control. However, gallic acid at all doses significantly reverted the observed reduction in erythrocyte and heart SOD activities of negative controls, with doses higher than 25 mg/kg body weight reverting it to the range of normal controls comparing favourably with ascorbic acid (Figure 4).

**Fig 4.**
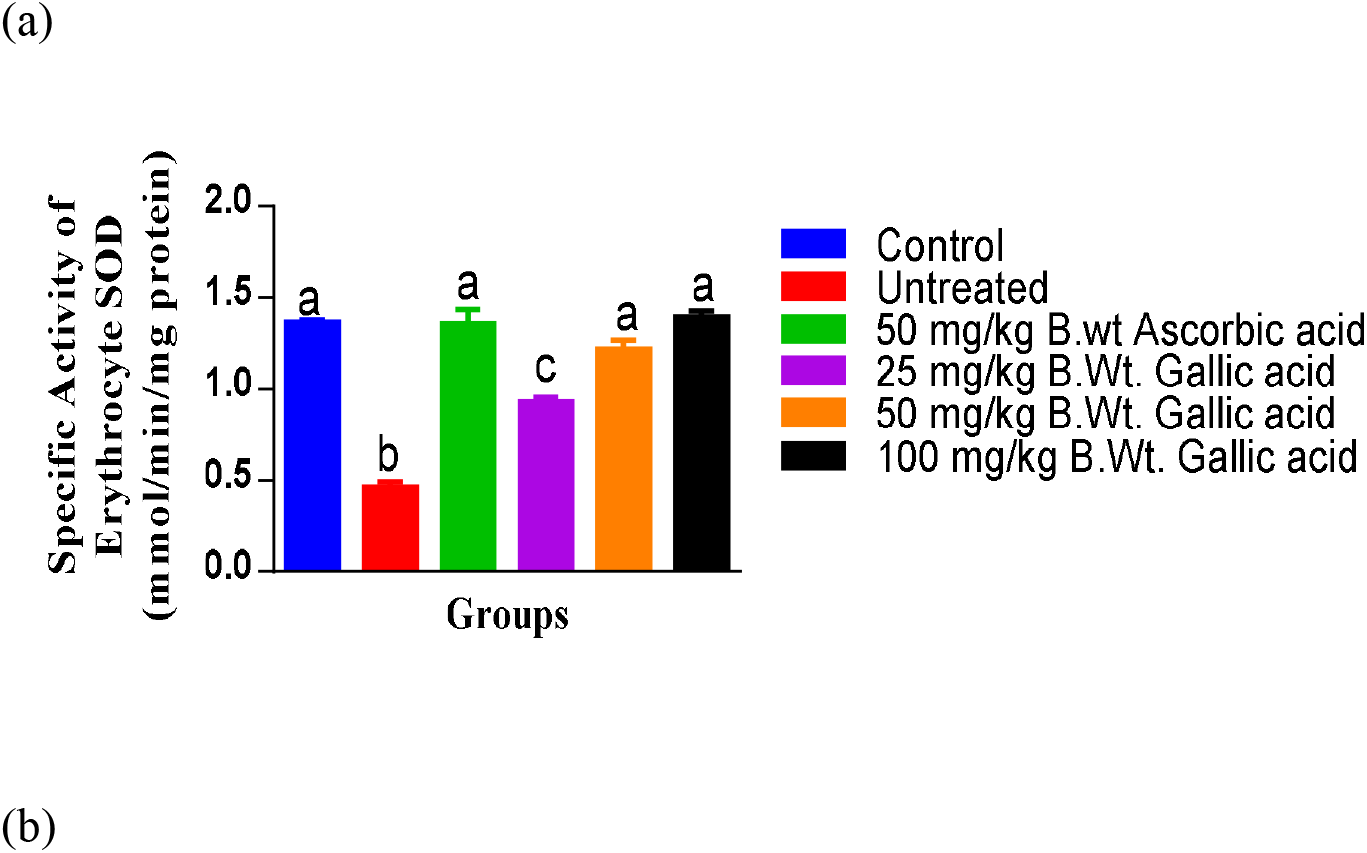
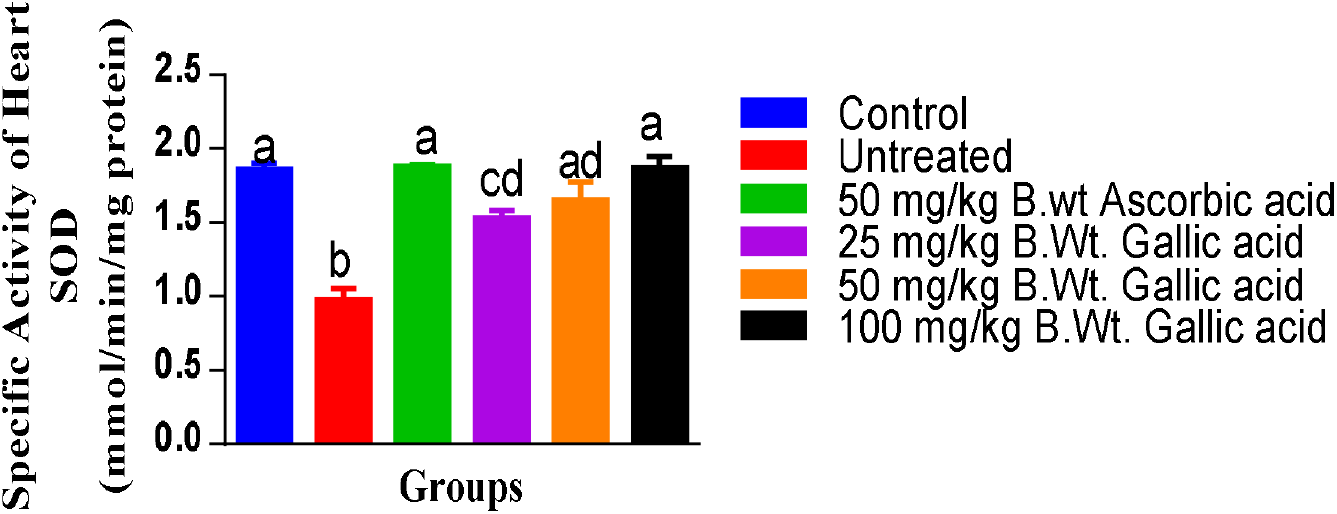
Effects of gallic acid on erythrocyte (a) and heart (b) superoxide dismutase (SOD) specific activity in mice with benzene-induced myelotoxicity. Values are means ± SEM of 6 replicates. Values for heart/erythrocyte with different superscripts are significantly different at p<0.05.

### Catalase (CAT)

There was significant reduction in erythrocyte and heart CAT activities of mice with benzene-induced myelotoxicity (negative controls) compared to normal controls. Gallic acid at all doses significantly reverted the observed reduction in erythrocyte and heart CAT activities of negative controls to range of normal controls, comparing favourably with ascorbic acid (Figure 5).

**Fig 5.**
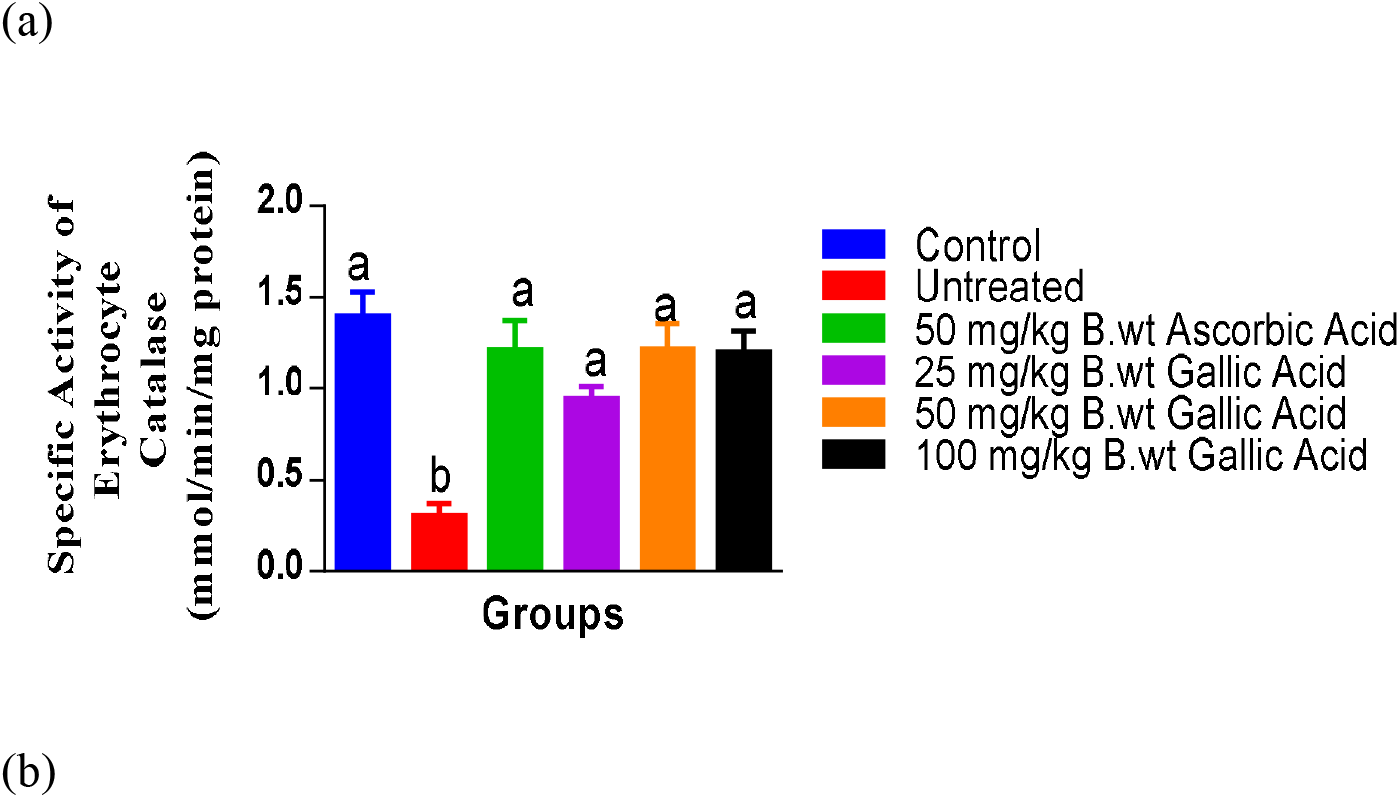
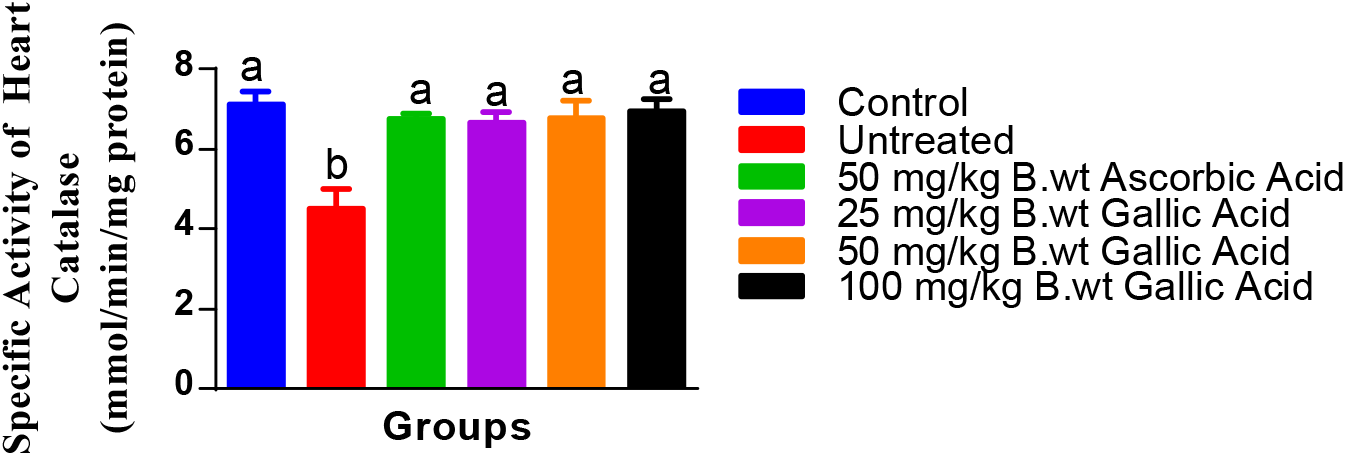
Effects of gallic acid on erythrocyte (a) and heart (b) catalase (CAT) specific activity of mice with benzene-induced myelotoxicity. Values are means ± SEM of 6 replicates. Values for heart/erythrocyte with different superscripts are significantly different at p<0.05.

### Glutathione peroxidase (GPx)

There was significant reduction in erythrocyte and heart GPx activities of mice with benzene-induced myelotoxicity (negative controls) compared to normal controls. Gallic acid at all doses significantly reverted the observed reduction in erythrocyte and heart GPx activities of negative control to the range of normal control, comparing favourably well with ascorbic acid (Figure 6).

**Fig 6.**
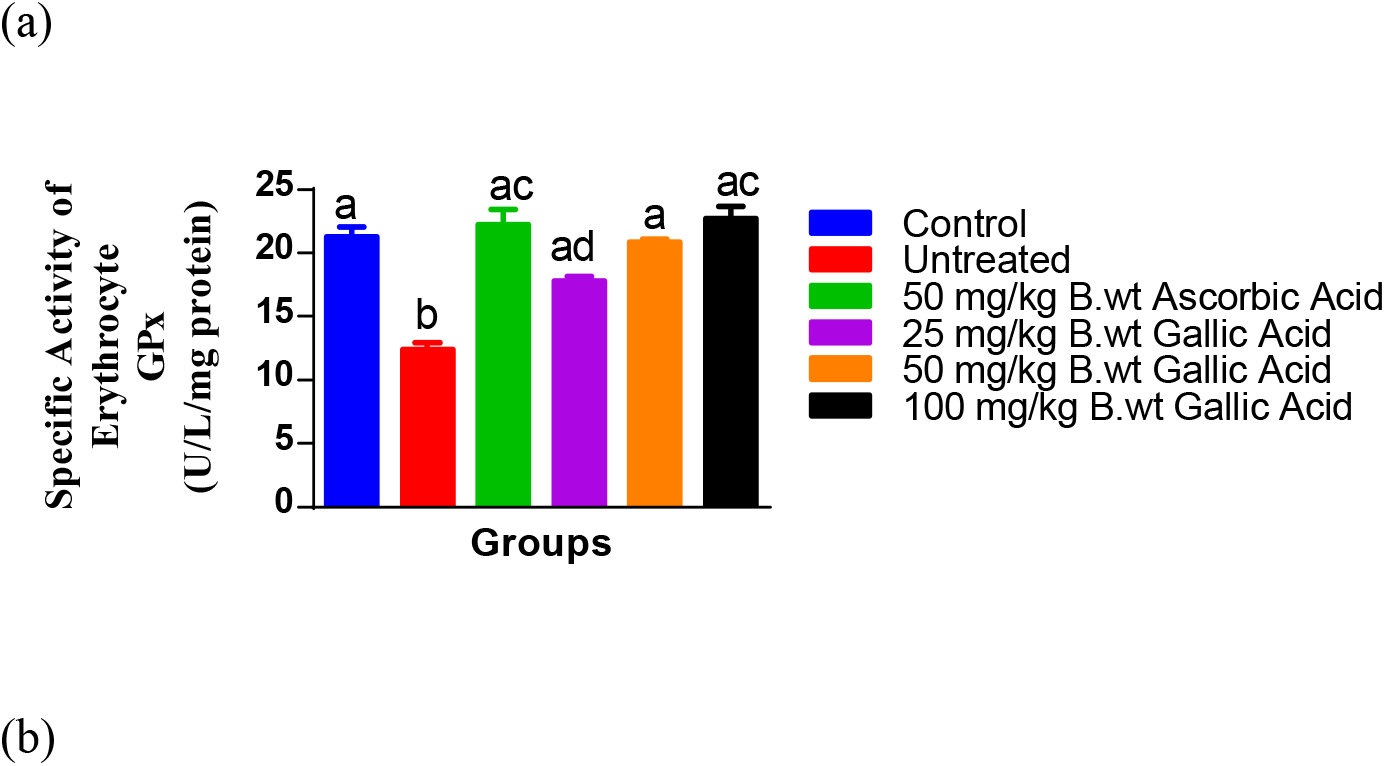
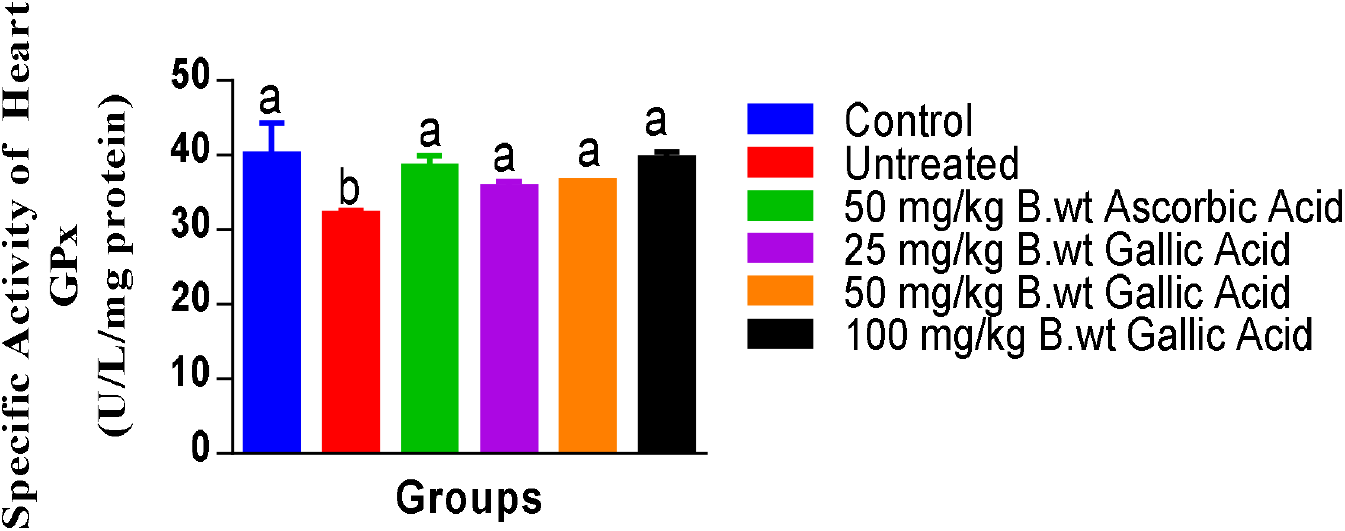
Effects of gallic acid on erythrocyte (a) and heart (b) glutathione peroxidase (GPx) specific activity of mice with benzene-induced myelotoxicity. Values are means ± SEM of 6 replicates. Values for heart/erythrocyte with different superscripts are significantly different at p<0.05.

### Reduced glutathione (GSH)

There was significant reduction in erythrocyte and heart GSH concentrations of mice with benzene-induced myelotoxicity (negative controls) compared to normal controls. Gallic acid at all doses significantly reverted the observed reduction in erythrocyte and heart GSH concentrations of negative controls to the range of normal controls, comparing favourably well with ascorbic acid (Figure 7).

**Figure 7.**
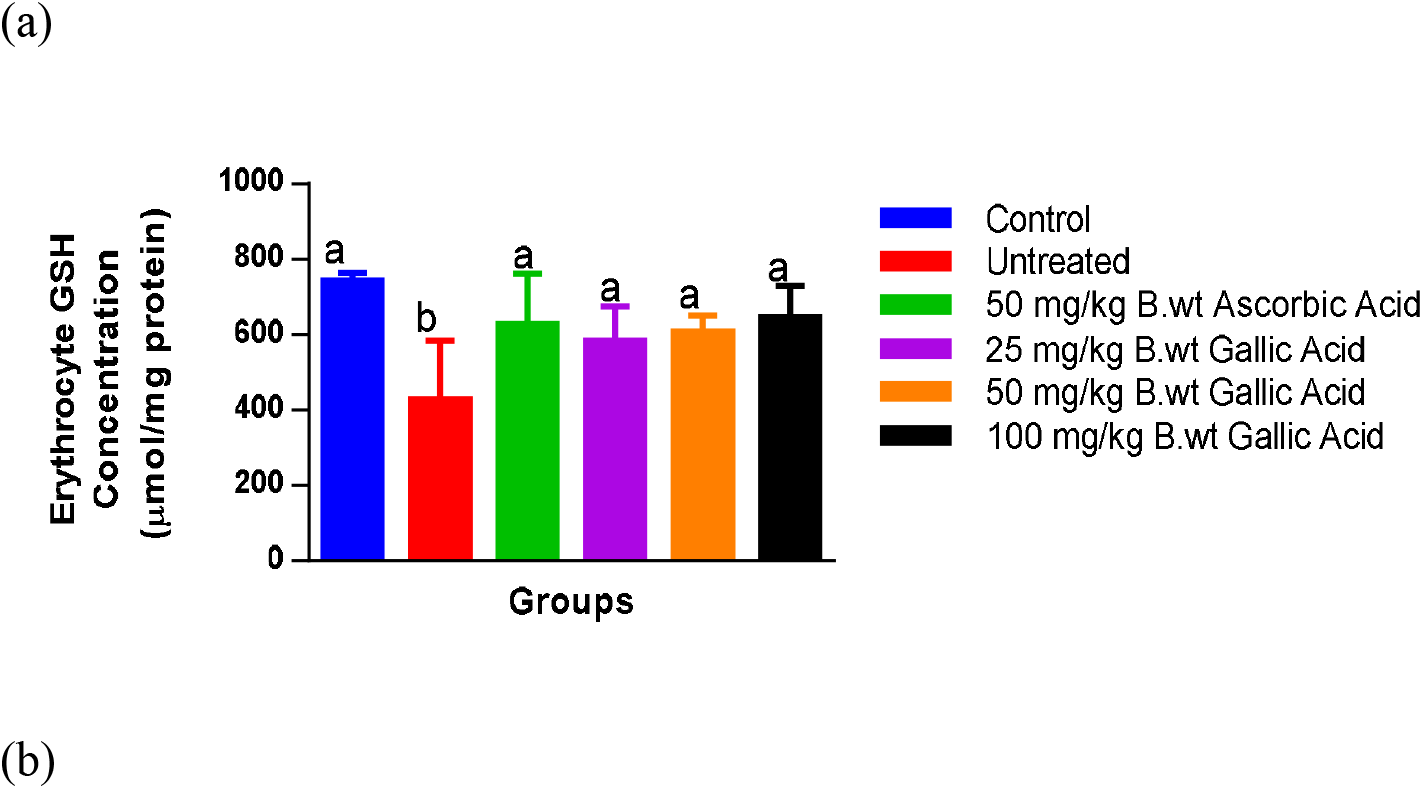
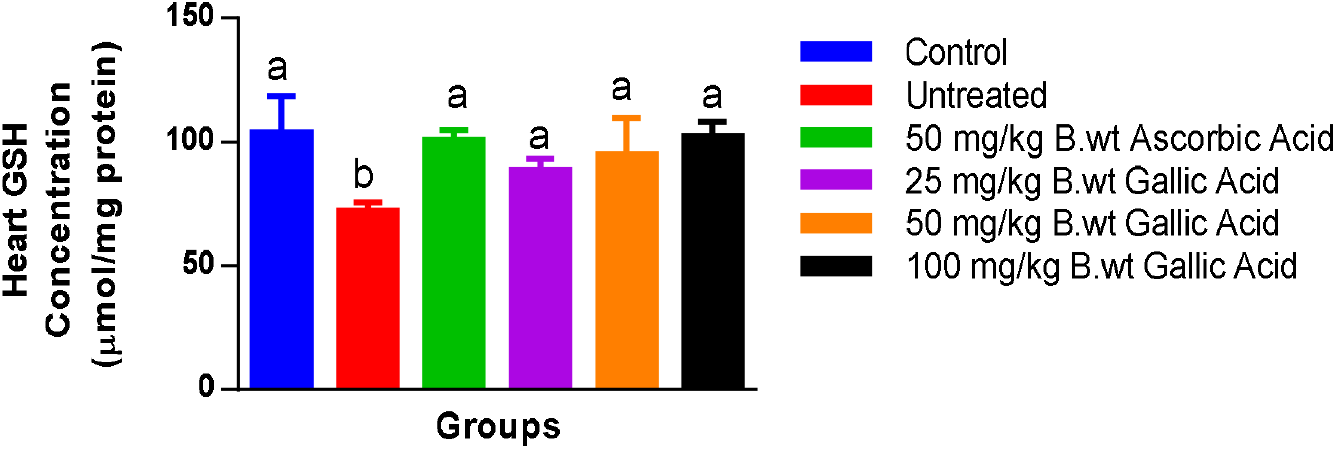
Effects of gallic acid on erythrocyte (a) and heart (b) reduced glutathione (GSH) concentration of mice with benzene-induced myelotoxicity. Values are means ± SEM of 6 replicates. Values for heart/erythrocyte with different superscripts are significantly different at p<0.05.

### Glutathione-S-transferase (GST)

Benzene caused significant reduction in erythrocyte GST activity of negative control compared to normal control. However, gallic acid at all doses significantly reverted the observed reduction in erythrocyte GST activity to the range of normal control, comparing favourably with ascorbic acid (Figure 8).

**Fig 8.**
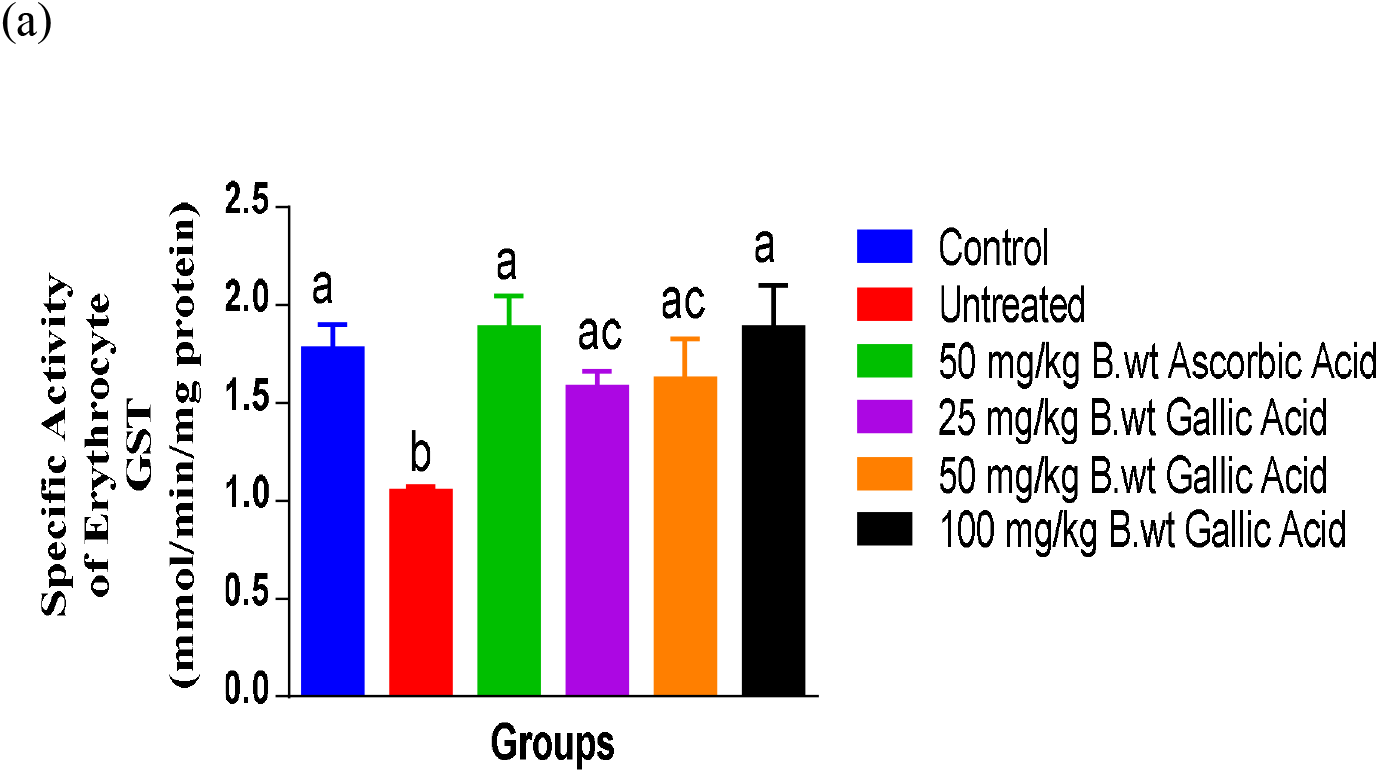
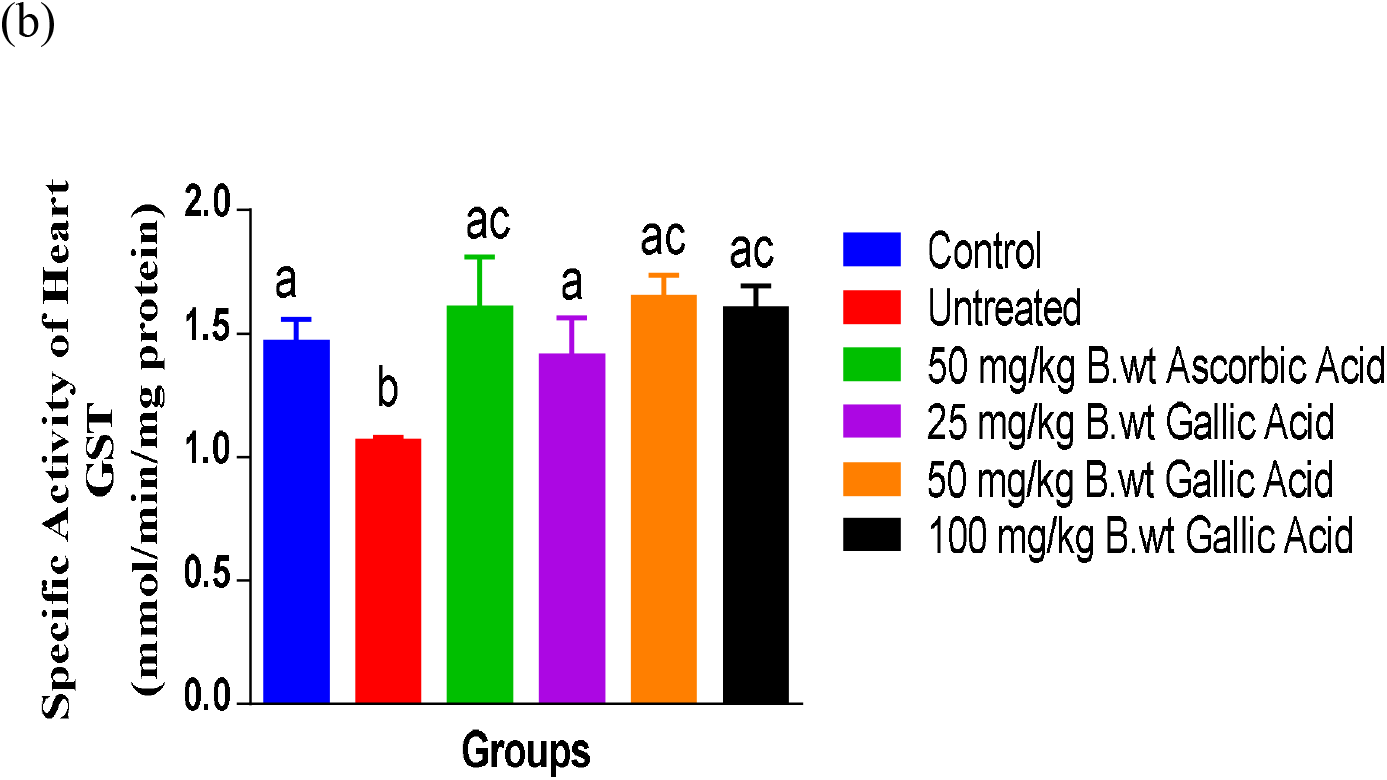
Effects of gallic acid on erythrocyte (a) and heart (b) glutathione-S-transferase (GST) specific activity of mice with benzene-induced myelotoxicity. Values are means ± SEM of 6 replicates. Values for heart/erythrocyte with different superscripts are significantly different at p<0.05.

### Protein concentration (PC)

There was significant reduction in erythrocyte and heart protein PC of mice with benzene-induced myelotoxicity (negative controls) compared to normal controls. Gallic acid at all doses significantly reverted the observed reduction in heart PC of negative control to the range of normal control while only doses of 50 and 100 mg/kg body weight significantly reverted the observed reduction in erythrocyte PC concentration of negative control to the range of normal control comparing favourably well with ascorbic acid (Figure 9).

**Figure 9.**
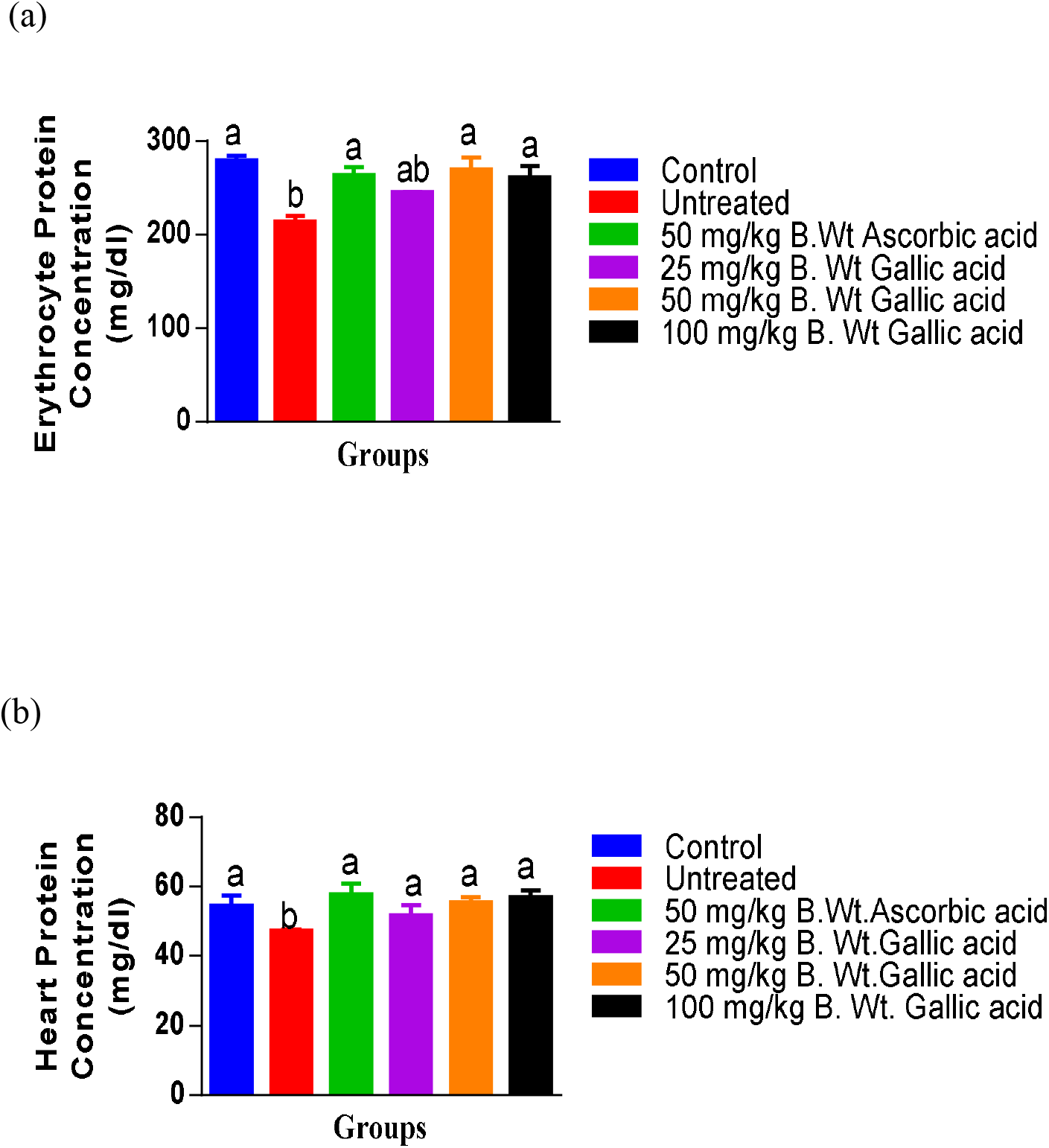
Effects of gallic acid on erythrocyte (a) and heart (b) protein concentration (PC) of mice with benzene-induced myelotoxicity. Values are means ± SEM of 6 replicates. Values for heart/erythrocyte with different superscripts are significantly different at p<0.05.

## Discussion

### The role of oxidative stress in benzene-induced myelotoxicity and secondary complications

Benzene-induced myelotoxicity usually occurs in humans as a result of occupational exposure to benzene (Cooper and Snyder, 1988; Yin *et al*., 1987). Several mechanisms have been proposed to explain benzene-induced myelotoxicity but, despite extensive research, the mechanism by which benzene induces myelotoxicity is still not clear. However, studies have shown that benzene metabolites are the primary causes of oxidative stress experienced during myelotoxicity (Rickert *et al*., 1979; Irons *et al*., 1982; Powley *et al*., 2000). Benzene is metabolized by cytochrome P450 2E1 (CYP2E1), one of the many isozymes of hepatic cytochrome P450 mixed function oxidases (Irons *et al*., 1982; Lovern *et al*., 1997; Powley *et al*., 2000). Benzene oxide, the first intermediate in CYP 2E1-mediated metabolism, is converted into a number of metabolites including phenol, hydroquinone, benzoquinone, catechol, and muconic acid/muconaldehyde. In the bone marrow, myeloperoxidase further oxidizes these phenolic metabolites of benzene to form free radicals capable of damaging the bone marrow (Sawahata *et al*., 1985; Kipen *et al*., 1988).

The harmful impact caused by toxic radicals in the bone marrow creates a pathway for the onset of many subsequent complications. For example, free radical suppression or damage to the myeloid progenitor cells of bone marrow impairs the production or reduce lifespan of erythrocytes, ultimately leading to varying degrees of anemia (Robert *et al*., 2000). In addition, chemical-induced hemolytic anemia is a condition in which exposure to certain chemicals such as benzene results in damage to the cytoskeletal membrane proteins and the eventual destruction of red blood cells (Robert *et al*., 2000). Benzene toxicity has also been linked to acute myeloid leukemia. Since mature differentiated blood cells arise from a pluripotent stem cell via sequential complex interactions of regulatory molecules and cellular controls, potential disruptions of these processes by oxidants may lead to pathogenesis of leukemias or lymphomas (Rinsky et al. 1987). Overall, oxidative stress has been implicated as either the primary cause of pathologies such as atherosclerosis, paraquat poisoning, radiation-induced pneumonitis and fibrosis, or as secondary contributor to disease progression including chronic obstructive pulmonary disease (COPD), cancers, type 2 diabetes, hypertension, cardiovascular and Alzheimer’s disease (Henry and Zhang, 2021).

### Effects of gallic acid on oxidative stress markers

It is well-known that free radical damage of biomolecules, accumulated under acute and chronic oxidative stress, is involved in the pathogenesis of wide spectrum of diseases (Kruk *et al*., 2019). In this study, the secondary cardiovascular complications of oxidative stress mediated by benzene in erythrocyte and heart of mice, and possible ameliorative effects of gallic acid were examined.

Results of selected oxidative stress biomarkers in erythrocyte and heart of negative control showed significant elevation in concentration of malondialdehyde (MDA), nitric oxide (NO), and protein carbonyl (PCO), as well as marked decrease in concentration of reduced glutathione (GSH) and protein compared to normal controls. These suggest that benzene by generating reactive metabolites caused lipid peroxidation and protein oxidation in erythrocyte and heart of mice (Stanely *et al*., 2009). Increased levels of NO, MDA, and PCO may imply that both reactive oxygen and nitrogen species substantially contributed to oxidative stress observed in target tissues. Furthermore, ^•^NO in the presence of elevated level of superoxide (O_2_ •^−^), one which often occurs as a result of impaired ability of superoxide dismutase (SOD), can generate peroxynitrite (ONOO^-^). Peroxynitrite in its protonated form (ONOOH) is a very strong oxidant that produces the reactive intermediates, nitrogen dioxide (^•^NO_2_) and hydroxyl radical (^•^OH). These oxidants rapidly oxidize macromolecules, including membrane lipids, structural proteins, enzymes, and nucleic acids (Henry and Zhang, 2021; Padma *et al*., 2013; Redler and Dokholyan 2012). Additionally, studies have reported deficiency of extracellular SOD leading to an increased level of superoxide, is associated with elevated blood pressure due to higher conversion of nitric oxide (a vasodilator) to peroxynitrite, ultimately serving as a risk factor for various cardiovascular diseases (Fattman *et al*., 2003). Meanwhile, oxidative stress resulting in the formation of lipid hydroperoxides, has been shown to underlie the conversion of LDL cholesterol into atherogenic form of oxidized-LDL (OxLDL), which has a crucial role in initiating and promoting the inflammatory response and recruitment of leukocytes in the lesion site, and contributes to the development of atherosclerosis through activation of smooth muscle cells and reduced NO bioavailability (Gniwotta *et al*., 1997). Also, elevated MDA and PCO levels have been linked to aberrant redox signalling of hydrogen peroxide (H_2_O_2_) that leads to accumulation of advanced glycation end products in diabetic complications (Evans *et al*., 2002; Castro *et al*., 2019; Liochev and Fridovich, 1994). Reduction in protein concentration may serve as an indication that, the integrity and function of cellular proteins is severely damaged due to oxidation or carbonylation, while that of GSH, being the most abundant non-enzymic antioxidant in the cell, has been compromised by depletion (Hsu and Yen, 2007).

However, administration of gallic acid considerably lowered NO, MDA, and PCO to levels similar to that of normal control. Also, concentrations of GSH and protein were increased with gallic acid treatment. These results demonstrate that gallic acid exhibited antioxidant effects, by scavenging reactive species excessively generated during oxidative stress that was facilitated by benzene metabolites in heart and erythrocyte of mice, thus corroborating earlier reports which indicated the roles of gallic acid in the prevention and treatment of oxidative stress complications in cancer, metabolic, cardiovascular, and neurodegenerative diseases, due to its ability to regulate redox homeostasis (Mileo and Miccadei, 2016; You and Park, 2010;Velderrain-Rodriguez *et al*., 2018; Mainzen *et al*., 2010).

### Effects of gallic acid on antioxidant enzymes

Results obtained from this experiment revealed marked reduction in the activities of catalase (CAT), glutathione peroxidase (GPx), glutathione-S-transferase (GST), and superoxide dismutase (SOD) in the heart and erythrocyte of mice with benzene-induced myelotoxicity in contrast to normal control. Here, inferences can be made that benzene, via its metabolites such as benzene oxide, phenol, hydroquinone, and benzoquinone, produced high amounts of free radicals that overwhelmed the capacity of antioxidant enzymes, the tissues first line of defense against oxidative damage. In oxidative stress, O_2_ ^•−^, H_2_O_2_, and ONOO^-^ can cause release of iron from iron-sulfur (Fe-S) clusters of metalloproteins, especially those involved in antioxidant defenses and redox reactions of electron transport in mitochondria, thereby inactivating them. Fe released can then bind with H_2_O_2_ (formed from one electron reduction of O_2_ ^•−^) in the so-called Fenton reaction to generate ^•^OH, an extraordinarily strong oxidants that will damage whatever molecule it is next to in cells (Henry and Zhang, 2021; You and Park, 2010). Also, maintaining redox homeostasis is important for cellular function. Studies have revealed that type 2 diabetes mellitus is a metabolic disease that arises as a result of aberrant signalling mediated by oxidants which led to disruptions in redox homeostasis (Evans *et al*., 2002).

Nevertheless, the administration of gallic acid to treatment groups significantly reverted the observed reduction in the activities of these antioxidant enzymes, suggesting that gallic acid may either protect them from oxidative damage or assist the enzymes with free radical scavenging roles. Gallic acid at doses of 50 and 100 mg/kg body weight elicited the best response to oxidative stress and compared favourably well with ascorbic acid (reference drug), making it as potent as ascorbic acid in protecting the erythrocyte and heart of mice from benzene toxicity. These findings support earlier studies that gallic acid could protect the cardiovascular system against formation of advanced glycation end products by inducing and maintaining the levels of antioxidant enzymes (El-Hussainy *et al*., 2016; Badhani *et al*., 2015; Umadevi *et al*., 2012; Shahrzad *et al*., 2001).

In summary, gallic acid is able to provide protection for cardiovascular system under benzene-induced oxidative stress. It can restore the series of enzymic and nonenzymic antioxidants, and attenuate cardiotoxic oxidants levels through free radical scavenging activity. Therefore, gallic acid could be developed as a versatile antioxidant with promising therapeutic and industrial applications.

## Conclusion

As oxidative stress is a component of many diseases, the development of effective antioxidant therapies still remains an important goal of research, one of which is the use of dietary supplements such as gallic acid. The results of this study highlight the significant ability of gallic acid to ameliorate oxidative stress and enhance the antioxidant defense system in the heart and erythrocyte of mice with benzene-induced myelotoxicity.

However, to tackle oxidative stress more effectively whether in disease pathogenesis or progression, research on antioxidant defenses and the use of exogenous antioxidants must be tailored or directed at preventing the formation of free radicals and reactive species like O_2_^•−^, ONOO^-, •^OH, H_2_O_2,_ and hypohalous acid (HOX), rather than scavenging them.

## Acknowledgements

The author thanks his supervisor with whom conversation, collaboration, and supervision concerning this publication has occurred.

## Author contributions

**Toba I. Olatoye:** Conceptualization, Writing-original draft and editing. **Adebayo J.O**.: Supervision, Writing-review and editing. Both authors were involved in researching data, discussion of content, writing the thesis and editing the manuscript before submission.

